# xOmicsShiny: an R shiny application for cross-omics data analysis and pathway mapping

**DOI:** 10.1101/2025.01.30.635740

**Authors:** Benbo Gao, Yu H. Sun, Xinmin Zhang, Tinchi Lin, Wei Li, Romi Admanit, Baohong Zhang

## Abstract

**Summary:** We developed xOmicsShiny, a feature-rich R Shiny-powered application that enables biologists to fully explore omics datasets across experiments and data types, with an emphasis on uncovering biological insights at the pathway level. The data merging feature ensures flexible exploration of cross-omics data, such as transcriptomics, proteomics, metabolomics, and lipidomics. The pathway mapping function covers a broad range of databases, including WikiPathways, Reactome, and KEGG pathways. In addition, xOmicsShiny offers several visualization options and analytical tasks for everyday omics data analysis, namely, PCA, Volcano plot, Venn Diagram, Heatmap, WGCNA, and advanced clustering analyses. The application employs customizable modules to perform various tasks, generating both interactive plots and publication-ready figures. This dynamic, modular design overcomes the issue of slow loading in R Shiny tools and allows it to be readily expanded by the research and developer community.

**Availability and implementation:** The R Shiny application is publicly available at: http://xOmicsShiny.bxgenomics.com. Researchers can upload their own data to the server or use the pre-loaded demo dataset. The source code, under MIT license is provided at https://github.com/interactivereport/xOmicsShiny for local installation. A full tutorial of the application is available at https://interactivereport.github.io/xOmicsShiny/tutorial/docs/index.html.

**Supplementary data:** Supplementary data are available at *bioRxiv* online.

## 1 Introduction

With the growing trend of applying high-throughput technologies in biological research, a large amount of data including RNA-Seq, proteomics, and metabolomics has been generated to tackle complex biological questions. Since each omics type represents only one aspect of the biological process, cross-comparison of different types of omics data provides a more comprehensive understanding of how these changes converge at the pathway level. Indeed, multi-omics approaches have been widely used in many research areas (Hasin, et al., 2017), such as developmental biology (Rabinowitz, et al., 2017; Talman, et al., 2018), cancer (Li, et al., 2023; Yuan, et al., 2022), metabolic processes (Williams, et al., 2022), and complex diseases such as Alzheimer’s disease (Iturria-Medina, et al., 2022; Kodam, et al., 2023). Furthermore, the multi-omics strategy bolsters the evidence of robust changes under certain conditions, which also facilitates biomarker discovery for diagnostics and therapeutics (Ghosh, et al., 2016; Ren, et al., 2016; Sun, et al., 2019).

Multi-omics datasets are typically processed and summarized into large tables to store quantitative measurements and differential expression analysis results. To gain further insights into the underlying biological changes at the systems biology level, researchers have developed tools to accommodate this challenge, such as MONGKIE (Jang, et al., 2016), WIlsON (Schultheis, et al., 2019), Argonaut (Brademan, et al., 2020), multiSLIDE (Ghosh, et al., 2021), OmicsAnalyst (Zhou, et al., 2021), MiBiOmics (Zoppi, et al., 2021), PaintOmics 4 (Liu, et al., 2022), etc. However, considering the amount and complexity of modern omics data modalities, these tools only partially address the bottleneck of data visualization and interpretation. For instance, flexible integration of cross-omics data remains challenging, and a comprehensive pathway mapping function is in demand.

To fill these gaps, we developed xOmicsShiny, a user-friendly R Shiny application for interactive analysis of multi-omics datasets. It is designed to advance data exploration and network/pathway-level interpretation for either single-omics or multi-omics data. In particular, it features integrative analysis by merging any gene/protein-related comparisons (such as RNA-Seq, proteomics) or compound-focused datasets (such as metabolomics, lipidomics). Users can navigate WGCNA or PCSF network analysis results and map the data to multiple pathway databases such as WikiPathways, Reactome, and KEGG. xOmicsShiny is powered by multiple state-of-the-art R packages and users have the flexibility to upload, store, and analyze their data using our public server or on their local computers.

## 2 Methods

### Overall Architecture

xOmicsShiny is a web application written using the Shiny package (http://shiny.rstudio.com/), in R programming language. R Shiny applications can be easily configured on a local computer or deployed on a public or private server, enhancing user accessibility and data privacy. To ensure efficient loading of multiple functions, we employed a modular design and created 14 modules (Fig. 1) for diverse analyses. Users only need to load the required modules when using them, which saves memory and time during application loading.

**Fig 1.**
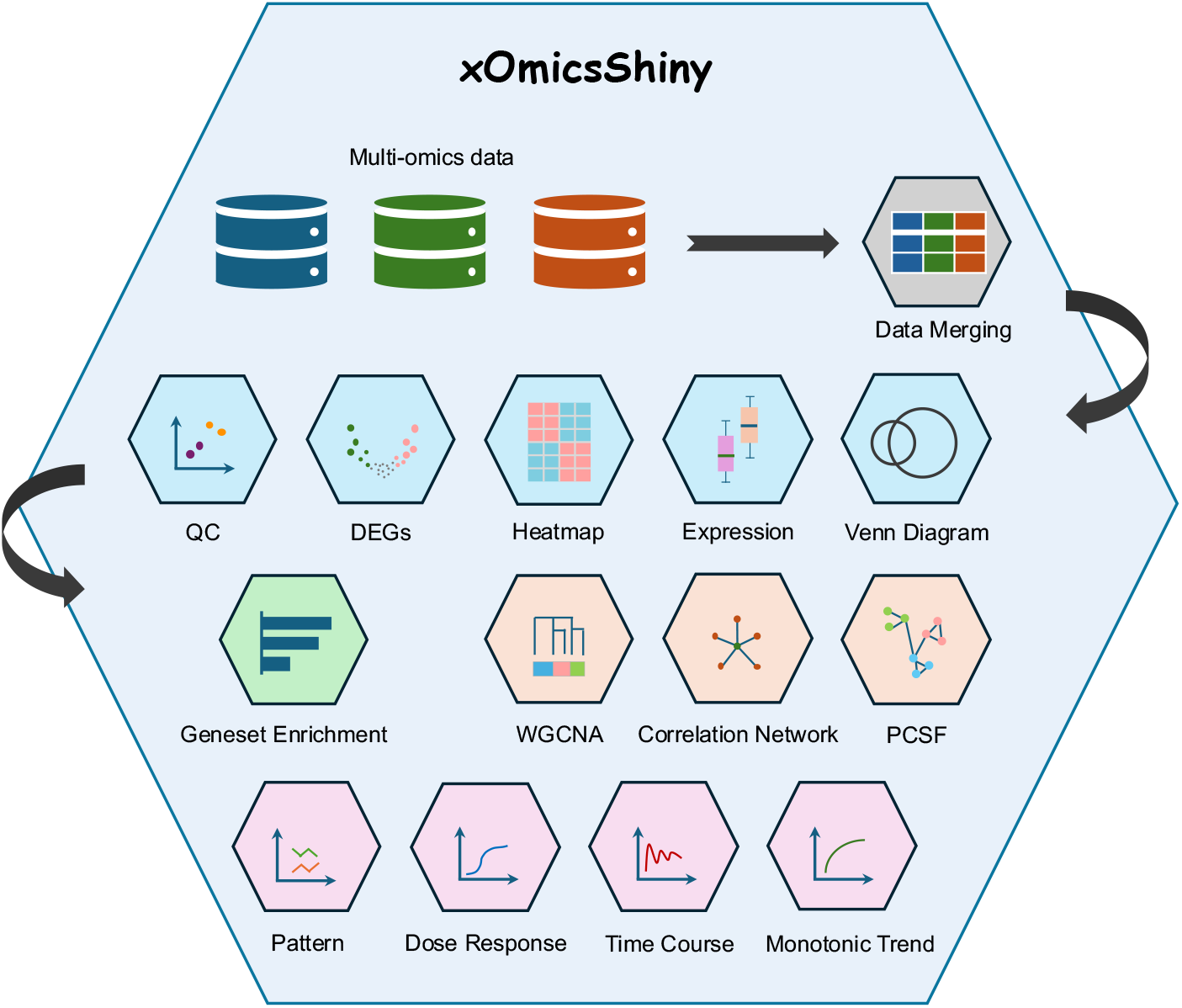
Overview of xOmicsShiny modules. The Shiny application handles multi-omics data including RNA-Seq, proteomics, and metabolomics data. The **Data Merging** (Grey) module integrates multi-omics data for downstream analysis. **Data Exploration** modules (Blue) contain QC, DEGs, Heatmap, Expression, and Venn Diagram modules. **Geneset Enrichment** (Green) module performs GSEA and pathway analysis. **Network Analysis** (Bisque) modules include WGCNA, Correlation Network, and PCSF modules. **Pattern and Trend Analysis** (Orchid) modules consist of Pattern, Dose Response, Time Course, and Monotonic Trend modules.

Following the best practices of bioinformatic analysis, we incorporated packages that have received broad recognition from the research community, including ComplexHeatmap (Gu, 2022; Gu, et al., 2016), fgsea (Korotkevich, et al., 2021), WGCNA (Langfelder and Horvath, 2008), PCSF (Akhmedov, et al., 2017), pathview (Luo and Brouwer, 2013), etc.

### Data Merging and Meta p-value Calculation

Merging data is a critical step in multi-omics analysis. xOmicsShiny automatically matches gene symbols and compound IDs and joins the differential expression results derived from multiple comparisons. While keeping the original information after merging, additionally, we provide four methods for meta p-value calculation: fisher, minP, simes, and stouffer. The meta fold change is displayed as the average fold change by default, but users can also provide a customized formula to summarize the fold changes.

### Enrichment Analysis and Pathway Visualization

As a key step in pathway analysis and prioritization, both Gene Set Enrichment Analysis (GSEA) and Over-Representation Analysis (ORA) are used in the ‘Geneset Enrichment’ module to calculate the enrichment statistics. The enrichment step is performed using the fgsea package (Korotkevich et al., 2016), and the visualization is facilitated by the collapsePathways function. We currently support WikiPathways, Reactome, and KEGG databases. A few examples of MetaBase pathways have also been included to demonstrate the versatile integration of commercially available databases into xOmicsShiny. The enrichment analysis is compatible with single-omics data (RNA-Seq, proteomics, or metabolomics data) or merged data containing both gene symbols and compound IDs. xOmicsShiny performs a one-step enrichment analysis using the KEGG database and displays both genes and compounds on the KEGG pathway map. To streamline this step, we created a metabolite database by combining multiple databases and constructing lookup tables across various ID formats. A total of 263,000 unique metabolites were extracted from PubChem, KEGG, HMDB, SMPDB, Metabolite RefMet, etc.

### Weighted Gene Co-expression Network Analysis (WGCNA)

We used the WGCNA R package to perform network analysis when multiple samples are available. The calculation may be time-consuming, so it will be run in the background and results will be saved. Users can easily load and explore the results once they are ready.

### Dose Response and Time Course Analysis

The Dose Response module offers multiple built-in models such as the log-logistic model, generalized log-logistic model, and Weibull model. The time course analysis automatically fits the data to several models such as linear, Gauss-probit, log Gauss-probit, as well as exponential models.

## 3 Features

xOmicsShiny is designed for comprehensive multi-omics data visualization, with an emphasis on understanding the biological mechanisms at the systems biology level. To this end, we developed 14 modules under the ‘Settings’ tab encompassing five major types of tasks: 1) Data merging, 2) Data exploration, 3) GSEA and pathway analysis, 4) Network analysis, and 5) Pattern and trend analysis. We have provided a detailed tutorial on how to process, upload, and navigate the data on xOmicsShiny. Required inputs should include original expression values, sample metadata, and differential expression statistics such as fold change, p-values, and adjusted p-values. Moreover, we curated public data containing RNA-Seq, proteomics, and metabolomics to serve as a demo dataset in the platform hosted by our public server. A set of supplementary figures featuring key functions of the application can be found in Supplementary data (Supplementary Fig. S1-11).

### Data Merging

The ‘Merge Data’ module is designed to merge multiple same or different types of omics data at the comparison level, rather than at the original data level. This can be achieved through a two-step procedure. First, users need to load multiple datasets they intend to merge. Then, under the ‘Merge Data’ module, users can select the desired comparisons and merge them together. The merged data will be displayed as a new dataset, containing meta fold change and p-values. Merging the same data type from different comparison groups highlights consistent changes across conditions, while cross-omics data integration facilitates pathway-level integrative analysis.

### Data Exploration

General data exploration consists of five modules: Quality Control (QC) module, Differentially Expressed Genes (DEGs) module, Heatmap module, Expression module, and Venn Diagram module. We leverage and improve several functions in the single-omics visualization platform, Quickomics (Gao, et al., 2021). The QC module generates various plots including principal component analysis (PCA) plot, sample-sample distance plot, dendrograms, etc. The DEGs module features volcano plot and scatter plot for visualization. For merged data, users can display side-by-side volcano plots or a merged volcano plot. Heatmap has been widely used in presenting high-throughput data, and the Heatmap module enables both static and interactive heatmap generation. Following that, the Expression module contains functions to display individual gene/compound values in various ways, such as boxplot, violin plot, etc. Lastly, users can examine overlapping genes using the Venn Diagram module. These modules are highly customizable, offering plenty of controls to adjust the output figures.

### GSEA and Pathway Analysis

The single ‘Geneset Enrichment’ module runs GSEA or ORA analysis on a variety of pathway and gene ontology databases, and maps multi-omics data to the pathway diagrams. For merged datasets, KEGG pathway maps can plot fold changes of multiple comparisons simultaneously. The enrichment analysis is performed onthe-fly as the users adjust fold change and p-value cutoffs to select genes or compounds. All enrichment statistics and gene lists can be easily saved for further investigation.

### Network Analysis

xOmicsShiny offers three types of network analysis, computed by the WGCNA module, the Correlation Network module, and the PCSF module. WGCNA has been widely used to identify co-expression regulatory networks and hub genes for mechanistic discovery. The WGCNA module displays the dendrogram of hierarchical clustering, which allows users to adjust parameters for tree cutting. Following that, corresponding gene clusters will be shown for further investigation. The Correlation Network module, powered by the visNetwork and networkD3 R packages, provides interactive correlation network visualization on genes or compounds based on user-defined cutoffs. Moreover, we incorporated the Prize-collecting Steiner Forest (PCSF) method to further highlight sub-networks and functional units (Akhmedov et al., 2017).

### Pattern and Trend Analysis

Identifying patterns and trends has been critical to certain tasks, including analyzing dose-response treatment data or time-course experiment outputs. To achieve this, we developed four modules: Pattern module, Curve Fitting Dose Response module, Curve Fitting Time Course module, and Monotonic Trend module. The Pattern module identifies clusters on a set of user-defined genes using soft clustering, k-means, or partition around medoid (PAM) strategies. The optimal number of clusters can be determined by methods including silhouette coefficients and Within-Cluster Sum of Squares (WSS) values. The next two modules, as their names suggest, perform dose response curve fitting and time course fitting analysis. The Monotonic Trend module is useful for identifying consistent trends in the data, especially when the changes are not linear.

## 4 Conclusions

In summary, xOmicsShiny provides a user-friendly solution to bridge the gap between high-dimensional, multi-omics data and biological insights, with high scalability and extensive features compared with other R Shiny applications (Gao, et al., 2021; Munk, et al., 2024; Tien et al., 2024; Zhao and Wang, 2022; Zoppi, et al., 2021) (Supplementary table S1, S2). Users can follow the detailed tutorial to compile their own data tables, which can be uploaded, stored, and analyzed by the Shiny application. xOmicsShiny contains 14 modules to perform various analyses, from general quality control and exploratory analysis to complex tasks and pathway-level visualization. This is ensured by many databases pre-loaded in the application, such as KEGG database, gene ontology database, Reactome pathway database, and WikiPathways, etc. Moreover, it generates high-resolution, publication-quality figures that biologists can use directly. xOmicsShiny is released as an open-source Shiny application, which facilitates the bioinformatics community to further enhance its functionalities.

## Conflict of Interest

B.G., Y.H.S., T.L., R.A. and B.Z. are employees of Biogen and hold stocks from the company. X.Z. and W.L. are employees of BioInfoRx, Inc.

## Notes

https://github.com/interactivereport/xOmicsShiny

### Summary of Updates

Revised the description of pathway enrichment analysis and data merging.

