## Supplementary data for "xOmicsShiny: an R shiny application for cross-omics data analysis and pathway mapping"

Benbo Gao[1,#], Yu H. Sun[1,#,*], Xinmin Zhang[2], Tinchi Lin[1], Wei Li[2], Romi Admanit[1], and Baohong Zhang[1,*]

[1]Research and Development, Biogen Inc., Cambridge, MA, USA.

[2]Data Science, BioInfoRx Inc., Madison, WI, USA.

[#]Contributed equally to the work. *To whom correspondence should be addressed.

Table of contents

Figure S1: Modular design of the main functions.

Figure S2: Cross-project analysis.

Figure S3: Venn diagram for cross-omics gene and protein comparison.

Figure S4: Data merging.

Figure S5: Gene expression plots with p-values highlighted.

Figure S6: Prize-collecting Steiner Forest (PCSF) network analysis.

Figure S7: Plots generated by the modules related to drug studies.

Figure S8: Geneset Enrichment module.

Figure S9: Pathway visualization.

Figure S10: Overlay of data on the WikiPathways diagrams.

Figure S11: Overlay of data on Metabase pathway diagrams.

Table S1: Running speed test of example datasets.

Table S2: Feature comparison with existing R Shiny applications.

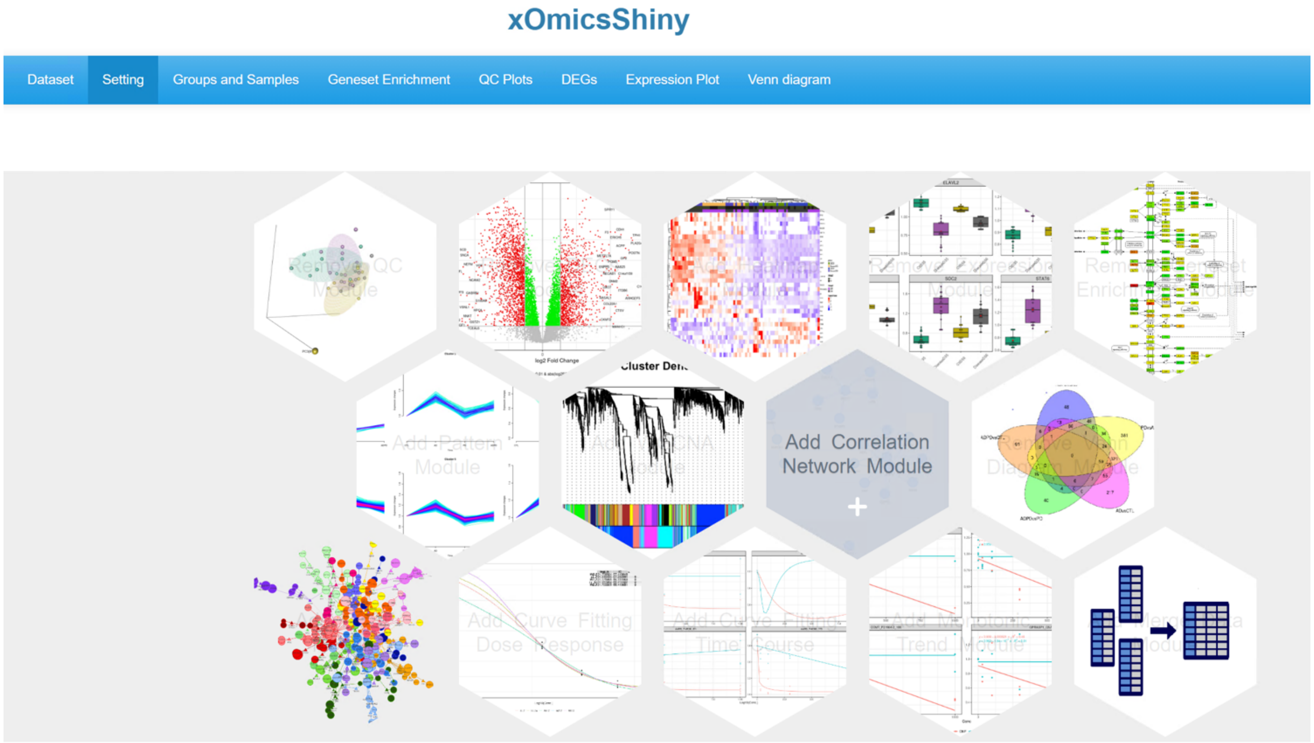


Figure S1: Modular design of the main functions. Users can mouse over a module to add or remove it. The ‘Correlation Network Module’ is selected in the screenshot.

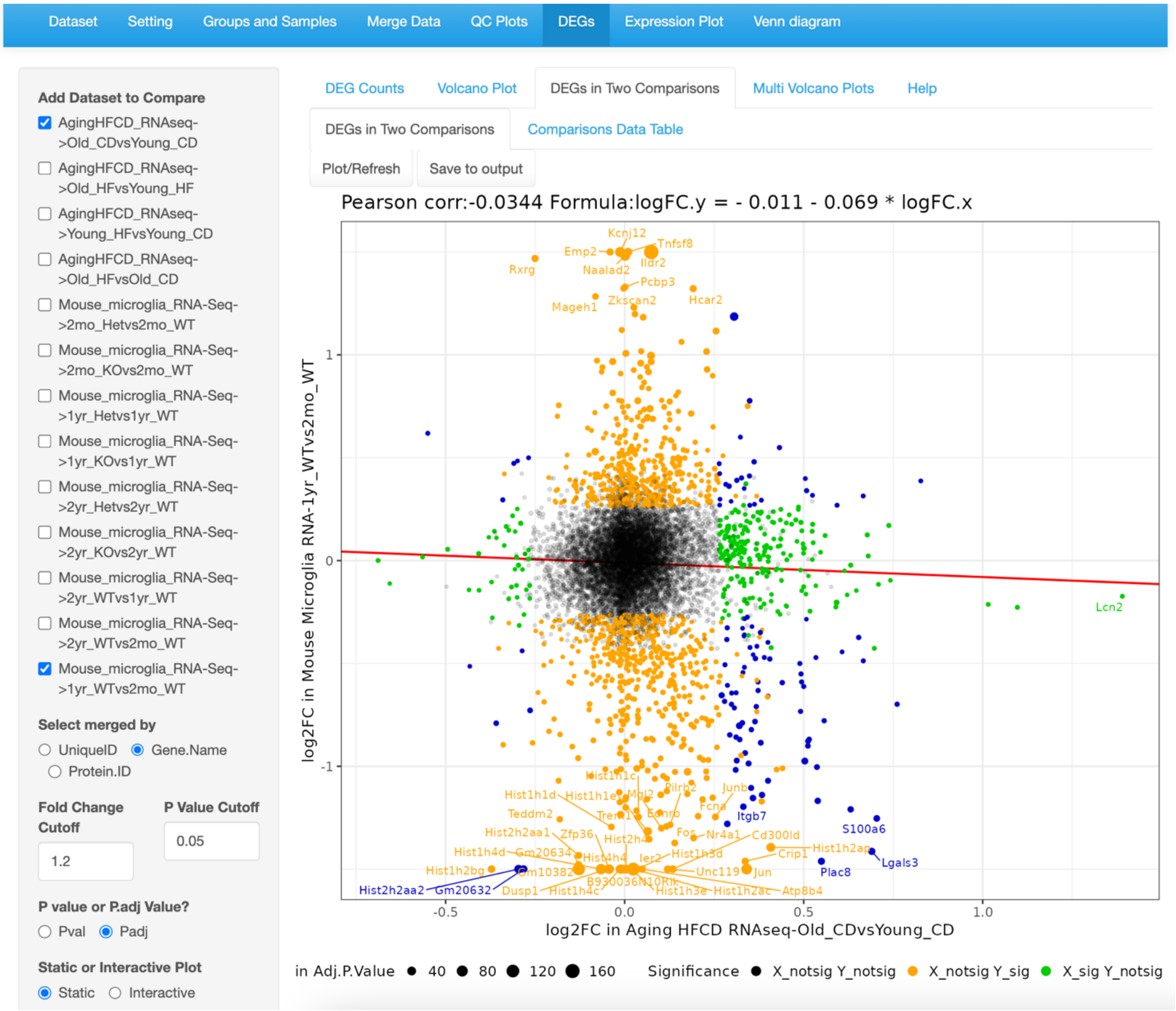


Figure S2: Cross-project analysis. Related comparisons from different studies can be compared directly to identify the common and distinct differentially expressed genes (DEGs).

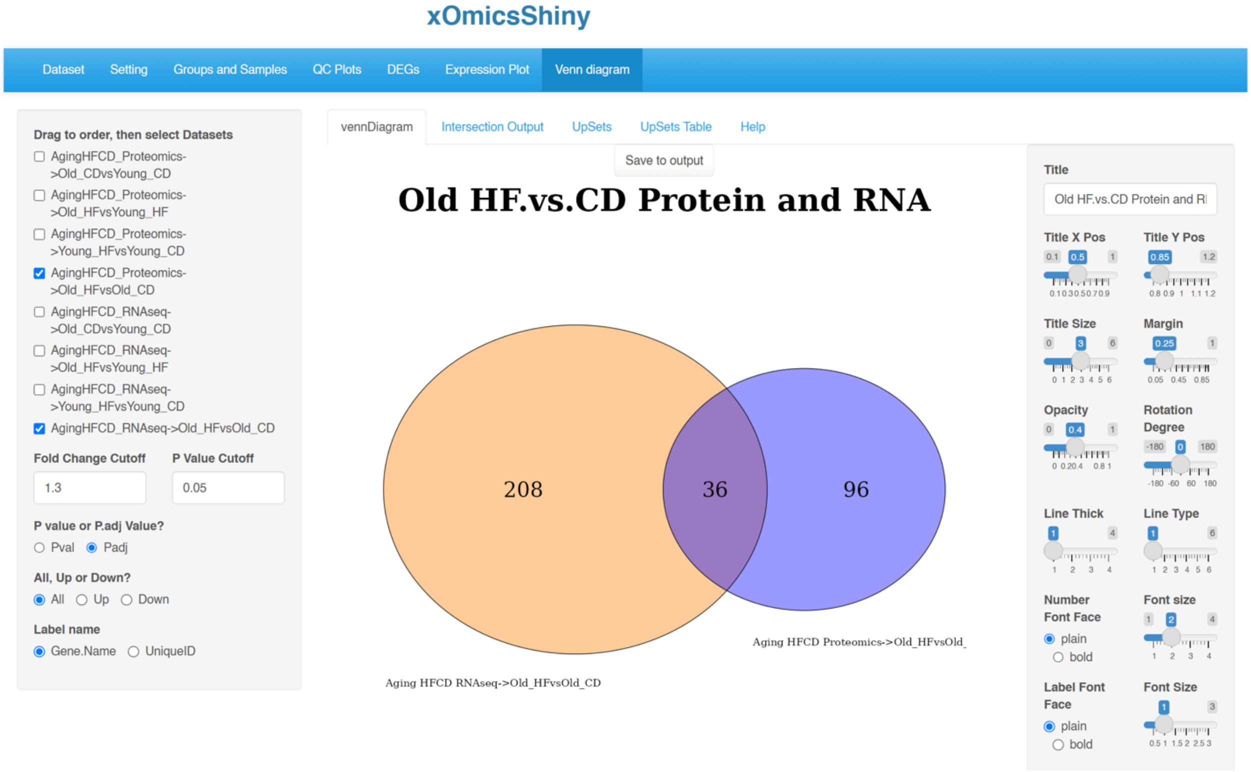


Figure S3: Venn diagram for cross-omics gene and protein comparison. The screenshot shows the overlapping significantly changed genes (left circle) and proteins (right circle) in the Old_HF (old mice, high fat diet treated) group compared with the Old_CD (old mice, control diet treated) group, in RNA-Seq and proteomics.

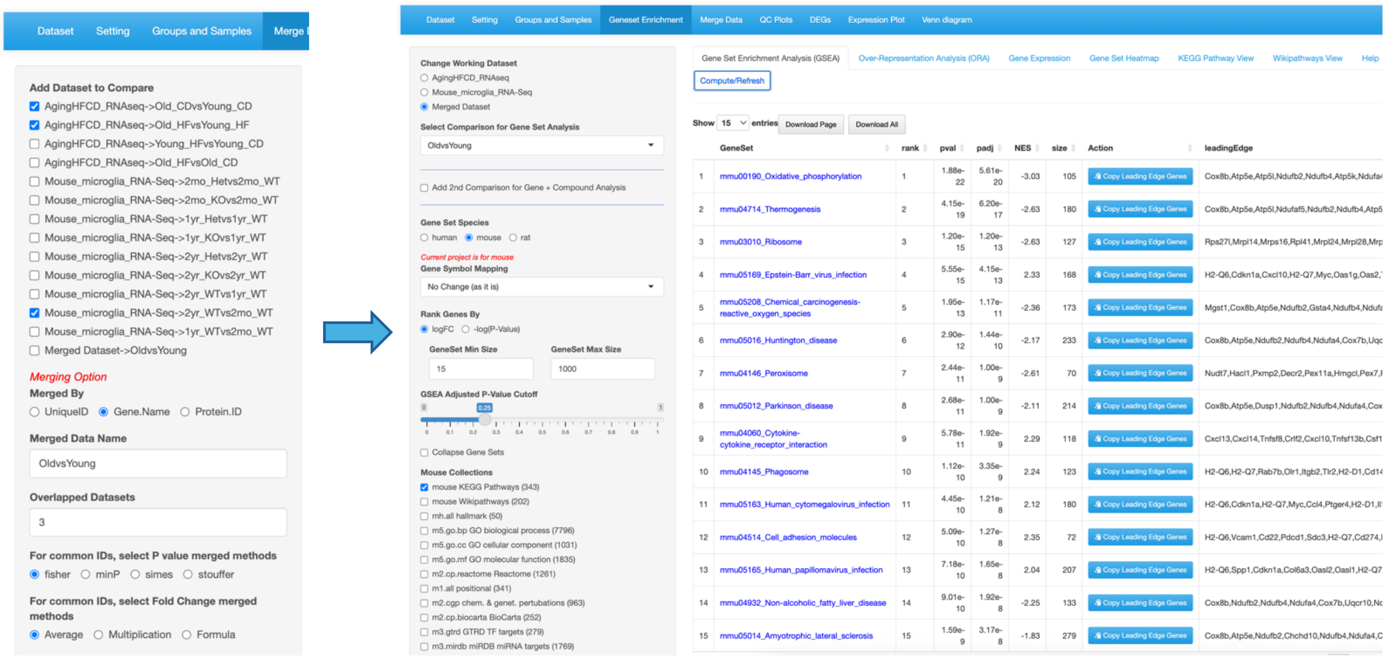


Figure S4: Data merging. Related comparisons from different studies (left panel) can be merged to retain common DEGs (right panel), which can be further explored using other modules, such as the Geneset Enrichment module.

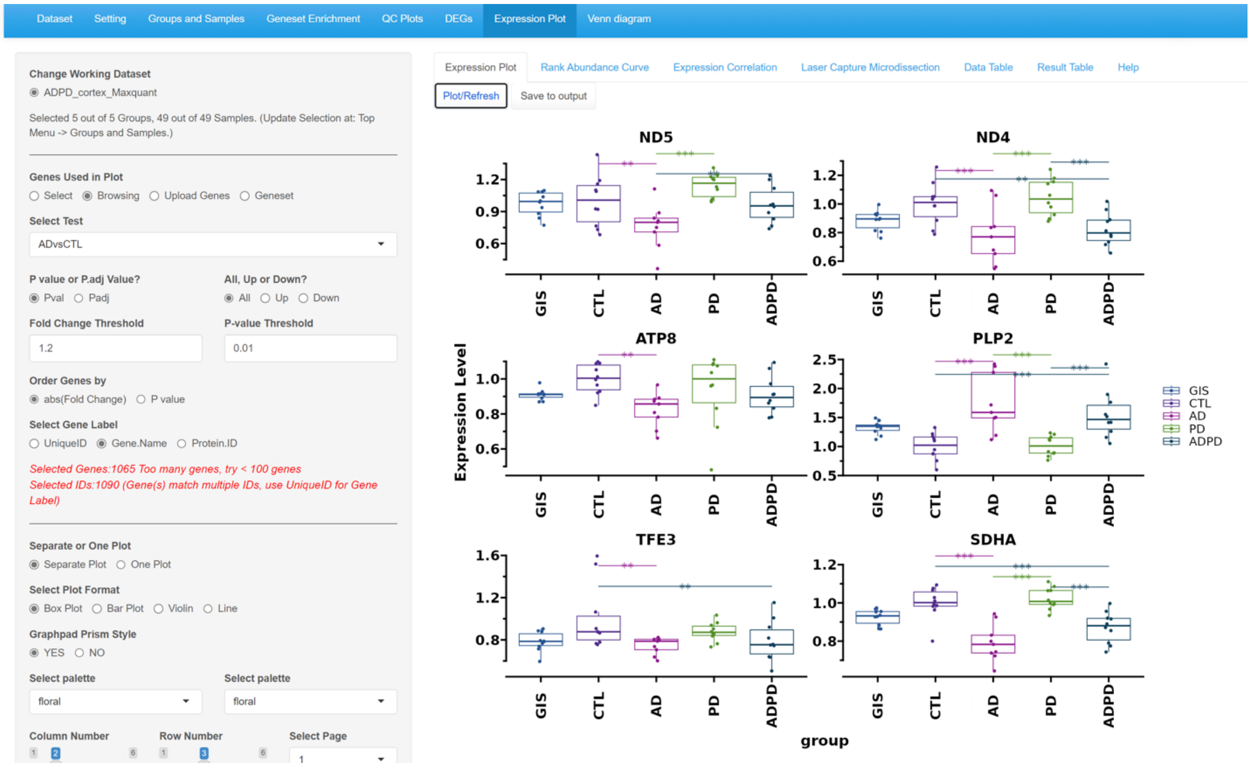


Figure S5: Gene expression plots with p-values highlighted.

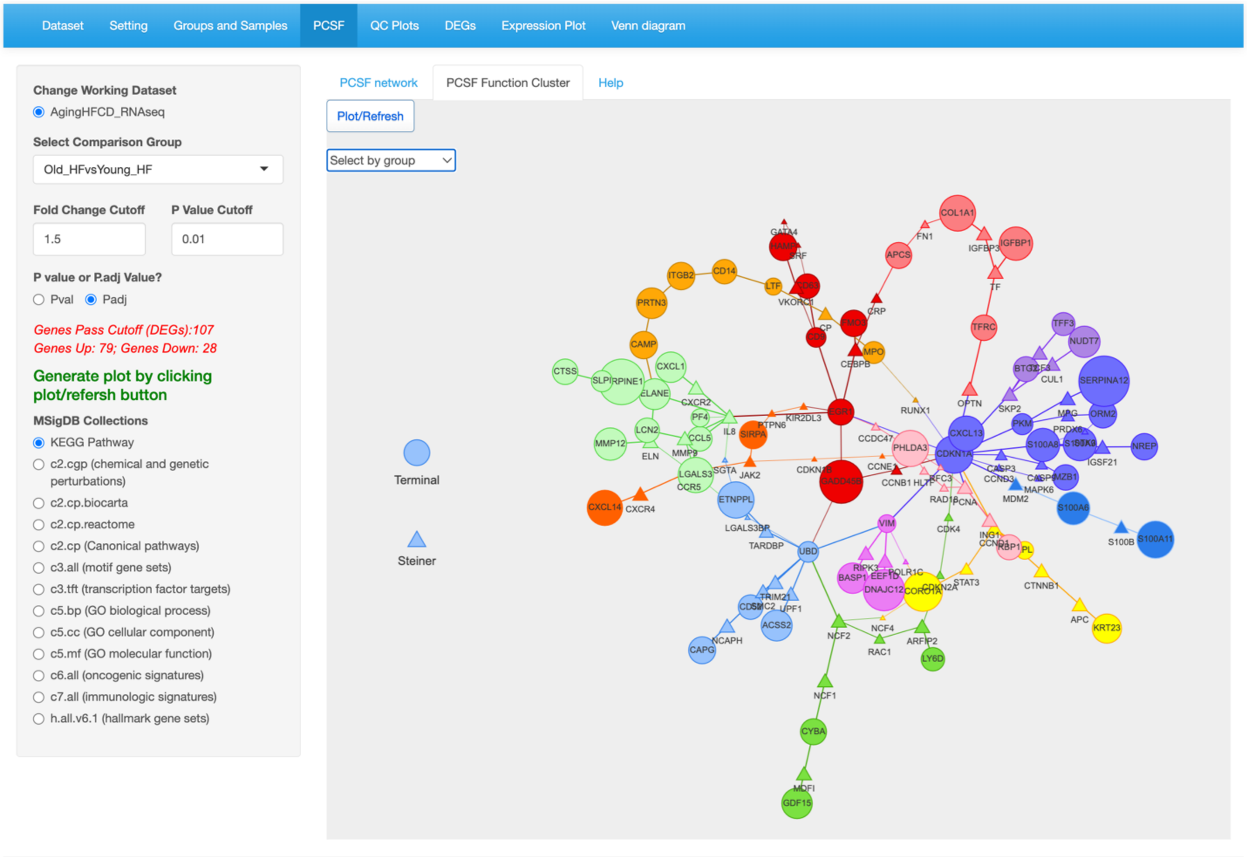


Figure S6: Prize-collecting Steiner Forest (PCSF) network analysis. The PCSF method performs mapping of the data to biological networks, such as protein-protein interactions, to gain further insights into the functional units.

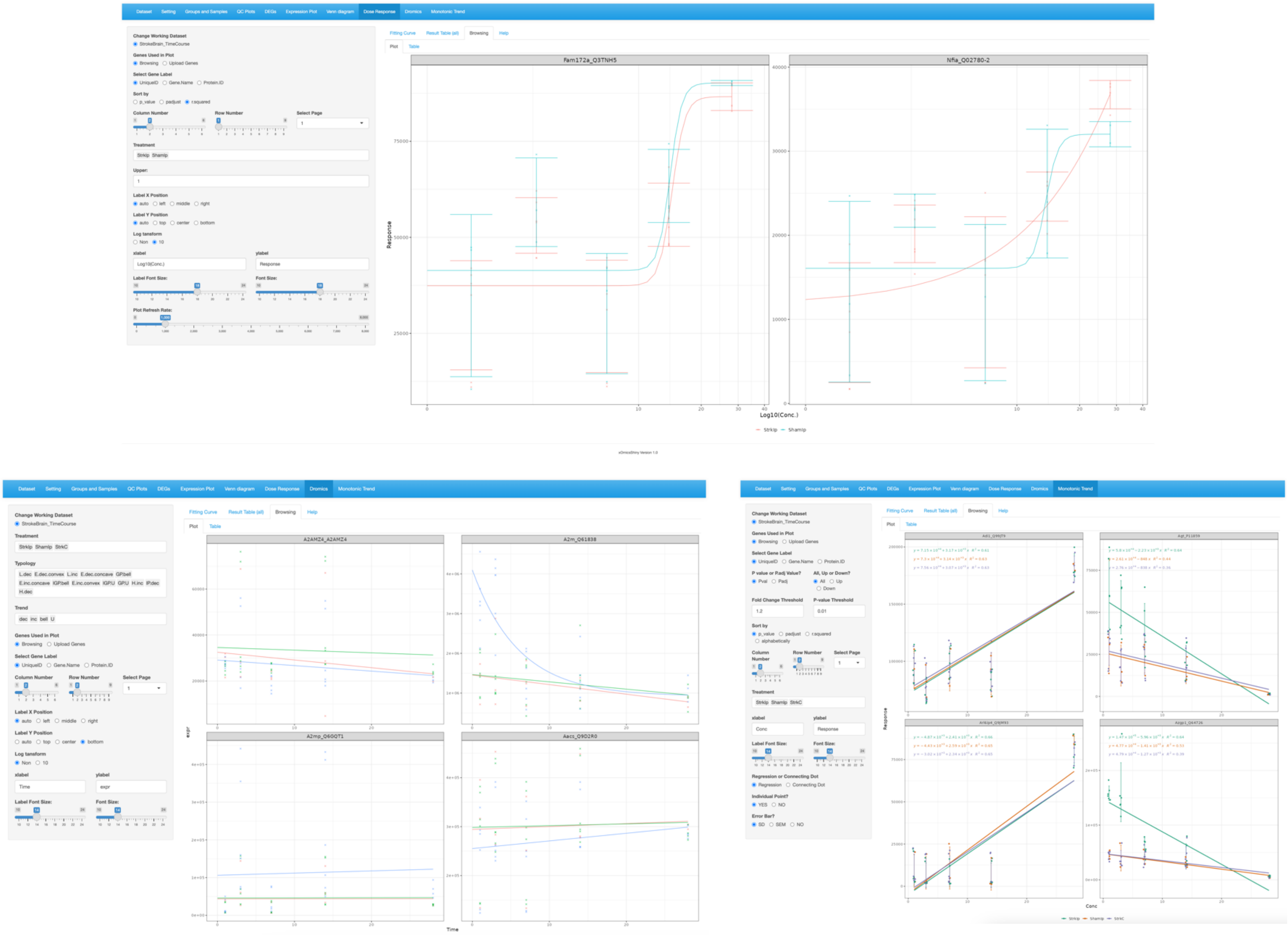


Figure S7: Plots generated by the modules related to drug studies. Dose Response (upper), Time Course (Dromics, lower left) and Monotonic Trend (lower right).

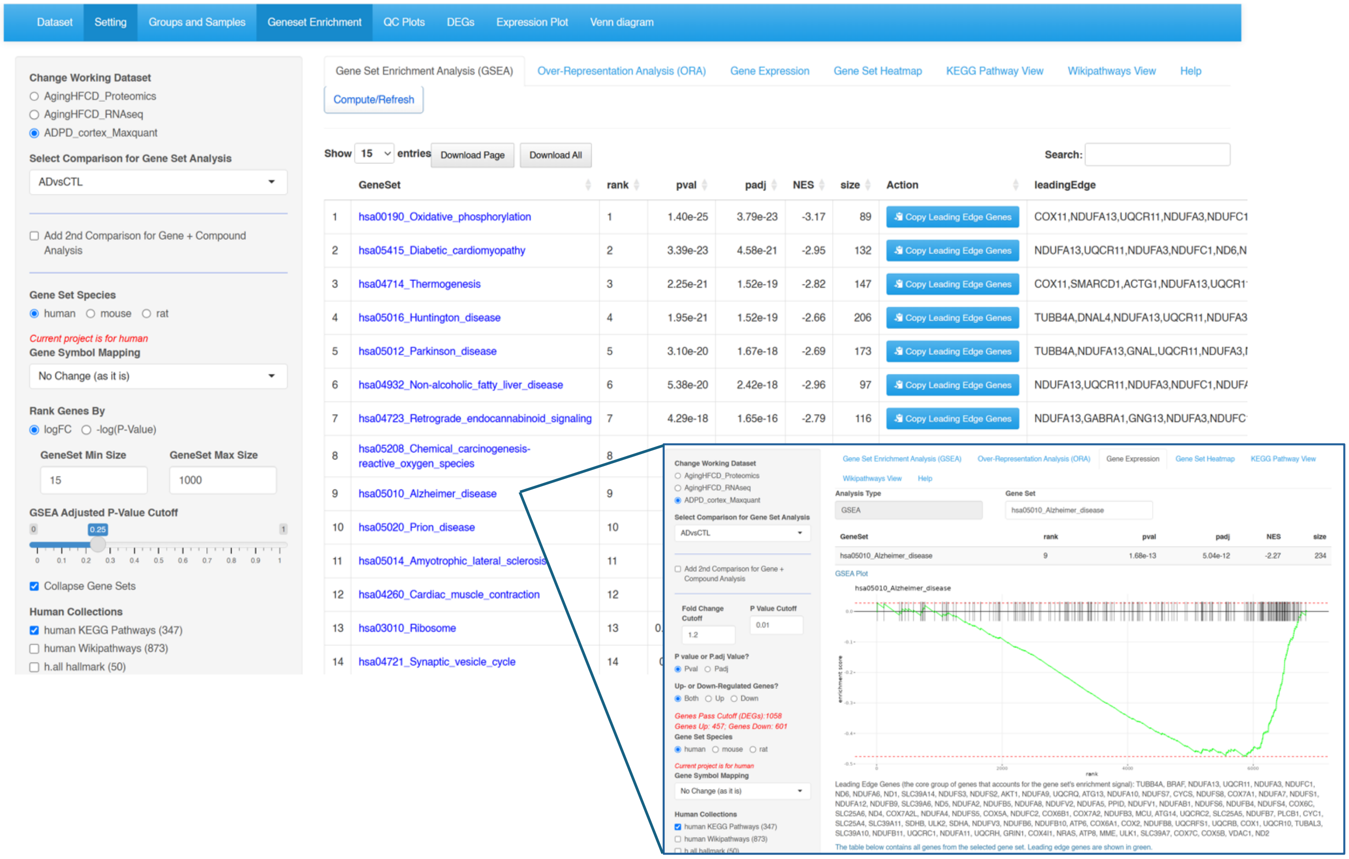


Figure S8: Geneset Enrichment module. Users can further explore the GSEA plots by clicking on the enriched pathway names (using the Alzheimer's Disease pathway as an example).

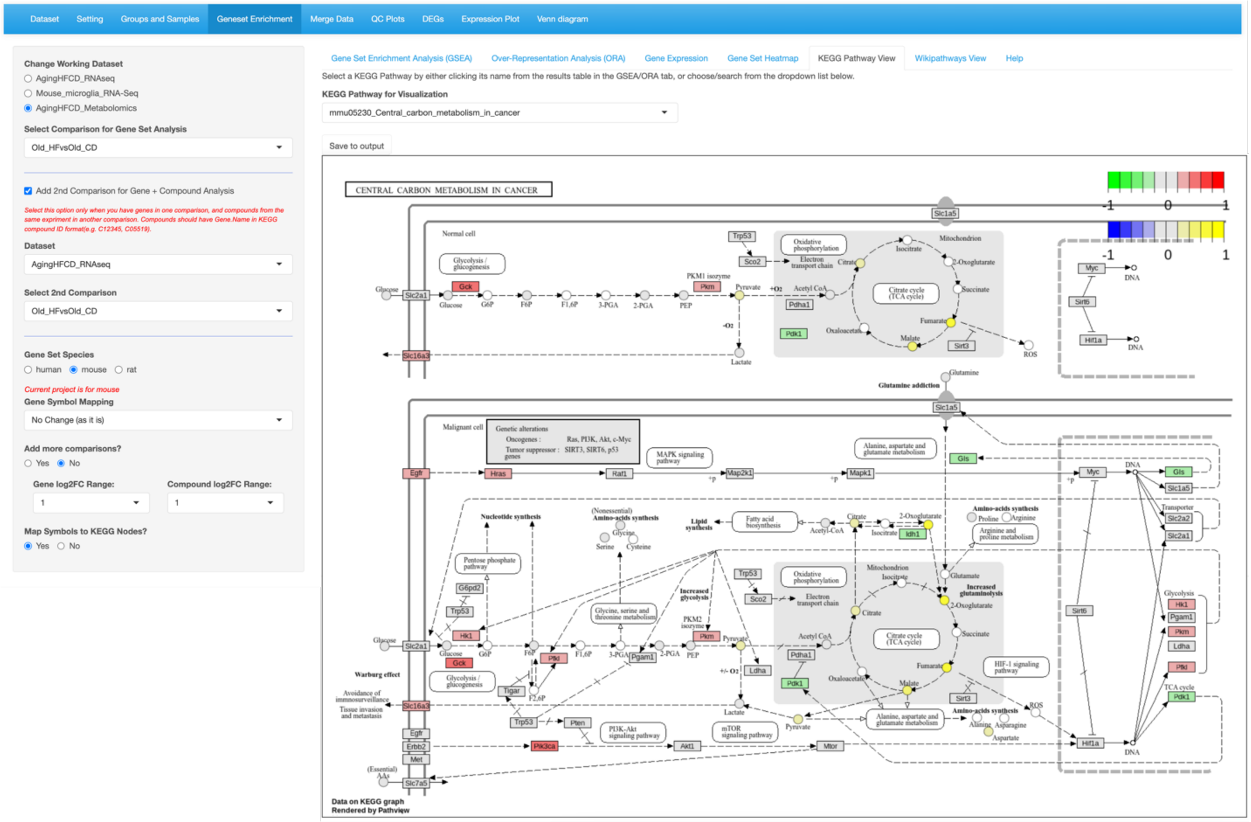


Figure S9: Pathway visualization by mapping genes and metabolites to KEGG pathway maps. Related comparisons from different studies can be merged and compared directly.

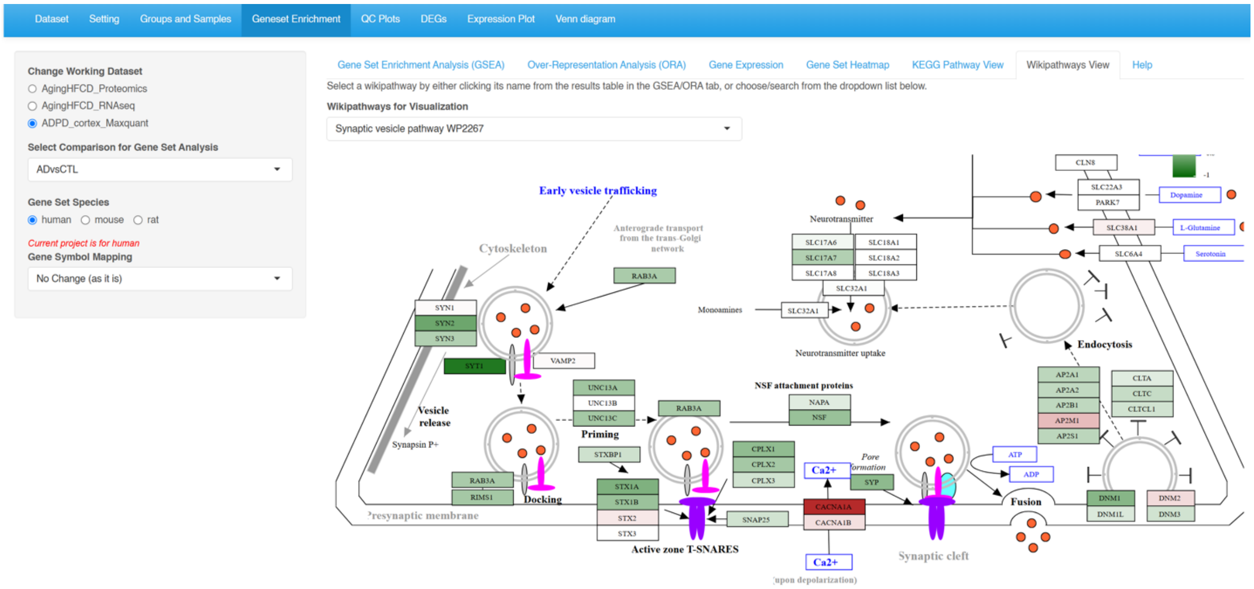


Figure S10: Overlay of data on the WikiPathways diagrams.

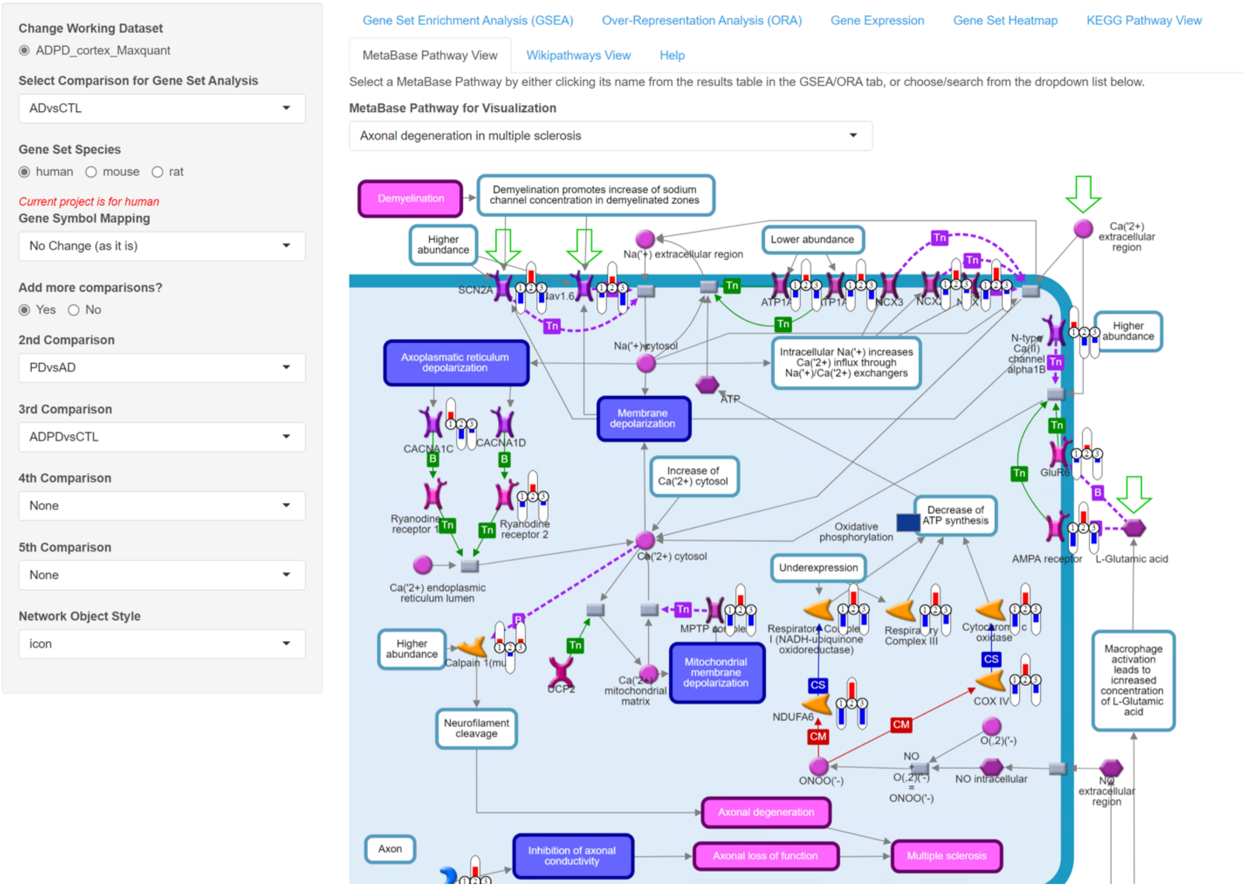


Figure S11: Overlay of data on Metabase pathway diagrams. This figure illustrates that the Metabase pathway maps are fully compatible with xOmicsShiny. Users may need to request a license to access the Metabase pathway database.

| Dataset | Samples | Comparisons | Measurements | Loading (s) | GSEA calculation (s) | | | | ORA calculation (s) | | | |
|---|---|---|---|---|---|---|---|---|---|---|---|---|
| | | | | | KEGG | GO, BP | GO, CC | GO, MF | KEGG | GO, BP | GO, CC | GO, MF |
| Mouse Microglia RNA | 93 | 9 | 15,402 | 9.2 | 3.8 | 17.1 | 8.1 | 8.1 | 1.3 | 15.7 | 2.7 | 3.7 |
| LRRK2 Neuron RNA | 54 | 2 | 15,168 | 7.1 | 7.8 | 24.3 | 6.4 | 10.2 | 1.7 | 19.1 | 3.3 | 4.7 |
| LRRK2 Neuron Proteome | 76 | 2 | 9,807 | 7.2 | 4.5 | 22.8 | 16.2 | 9.0 | 1.5 | 15.7 | 3.1 | 3.9 |
| ADPD cortex Searched by Maxquant | 49 | 6 | 7,163 | 5.3 | 2.9 | 7.8 | 4.4 | 4.2 | 1.0 | 12.1 | 2.4 | 3.1 |
| ADPD ACG Searched by Maxquant | 50 | 6 | 8,145 | 5.4 | 5.9 | 8.5 | 4.6 | 4.5 | 1.1 | 13.1 | 2.3 | 3.3 |
| ADPD cortex Searched by PD | 49 | 6 | 8,746 | 5.5 | 3.6 | 11.1 | 6.1 | 4.9 | 1.2 | 12.6 | 2.1 | 3.2 |
| ADPD ACG Searched by PD | 50 | 6 | 9,605 | 5.5 | 3.6 | 10.3 | 5.9 | 5.1 | 1.1 | 12.2 | 2.1 | 3.4 |
| Aging HFCD Metabolomics | 590 | 4 | 393 | 4.6 | 1.4 | NA | NA | NA | 1.1 | NA | NA | NA |
| Aging HFCD Proteomics | 315 | 4 | 3,941 | 6.5 | 3.3 | 7.2 | 4.1 | 3.9 | 1.1 | 12.1 | 2.1 | 3.1 |
| Aging HFCD RNAseq | 291 | 4 | 25,393 | 21.3 | 5.4 | 37.5 | 9.4 | 9.4 | 1.6 | 19.8 | 2.9 | 4.6 |
| Stroke Brain Time Course | 70 | NA | 7,565 | 7.1 | NA | NA | NA | NA | NA | NA | NA | NA |
| LCM 96Grid Simulated | 96 | 1 | 3,881 | 5.4 | 2.7 | 7.3 | 4.3 | 3.8 | 1.1 | 11.3 | 1.9 | 3.1 |

Table S1: Running speed test of example datasets. This table shows the number of samples, comparisons, and measurements (genes/proteins/metabolites) in each dataset. The additional columns record the loading speed (in seconds) of the data, as well as the time to calculate GSEA (Gene set enrichment analysis) and ORA (Over-representation analysis) results using the KEGG or gene ontology (GO) databases (BP: biological process, CC: cellular component, and MF: molecular function).

| Main features | | xOmicsShiny | Quickomics | Holomics | GraphBio | MiBiOmics | EasyPubPlot |
|---|---|---|---|---|---|---|---|
| Single-omics | General analysis | ✓ | ✓ | ✓ | ✓ | ✓ | ✓ |
| | PCA | ✓ | ✓ | ✓ | ✓ | ✓ | ✓ |
| | Volcano plot | ✓ | ✓ | | ✓ | | ✓ |
| | Heatmap | ✓ | ✓ | ✓ | ✓ | | ✓ |
| | Expression boxplot | ✓ | ✓ | | | | ✓ |
| | Venn Diagram | ✓ | ✓ | | | | |
| | Correlation analysis | ✓ | ✓ | ✓ | | ✓ | |
| | Network analysis (WGCNA, etc.) | ✓ | | | ✓ | ✓ | |
| | GSEA/GO analysis | ✓ | ✓ | | ✓ | | ✓ |
| | Pattern and trend analysis | ✓ | | | | | |
| Multi-omics | Multi-omics integration | ✓ | | ✓ | | ✓ | |
| | Multi-omics data exploration | ✓ | | ✓ | | ✓ | |
| | Multi-omics volcano plot | ✓ | | | | | |
| | Multi-omics pathway mapping | ✓ | | | | | |
| Citation | | Current work | Quickomics: exploring omics data in an intuitive, interactive and informative manner, Bioinformatics | Holomics-a user-friendly R shiny application for multi-omics data integration and analysis, BMC Bioinformatics | GraphBio: A shiny web app to easily perform popular visualization analysis for omics data, Frontiers in Genetics | MiBiOmics: an interactive web application for multi-omics data exploration and integration, BMC Bioinformatics | EasyPubPlot: a shiny web application for rapid omics data exploration and visualization, bioRxiv |

Table S2: Feature comparison with existing R Shiny applications.
